# Building a Tissue-unbiased Brain Template of Fibre Orientation Distribution and Tractography with Multimodal Registration

**DOI:** 10.1101/2022.04.13.488117

**Authors:** Jinglei Lv, Rui Zeng, Mai Phuong Ho, Arkiev D’Souza, Fernando Calamante

## Abstract

A brain template provides a standard space for statistical analysis of brain structure and function. For decades, the T1- and T2-weighted brain templates have been widely used for brain grey matter anatomical and functional analysis. However, T1- and T2-weighted templates provide very limited information about the axonal organization within the white matter. Recent advances in Diffusion MRI have enabled the detailed modelling of the axonal fibre orientation distribution (FOD) in white matter. Therefore, building a FOD template is essential for more robust white matter anatomy related analysis; however, it is important that this template aligns well with the cortical and subcortical structures. From such a FOD template, a tractography template can be also generated by fibre tracking algorithms, which can be used for subsequent applications, such as to perform the joint structural and functional analysis while ensuring rigorous fibre-to-fibre correspondence. In this paper, we explore the potential of generating the FOD template based on multimodal registration, in order to constrain the tempalte unbiased to both white and grey matter. We combine the information from T1-weighted, T2-weighted and Diffusion MRI to generate a coherent transformation for FOD registration and template generation. Our FOD template preserves the structural details at the white-grey matter boundary. To illustrate the benefit of this new approach, the resulting tractography template was used for joint structural-functional connectivity analysis.

## 1. Introduction

Diffusion MRI (dMRI) has attracted increasing attention in fundamental neuroscience(Assaf et al., 2019) as well as mental health research(Alexander et al., 2019; Calamante, 2019). For example, both the Human Connectome Project (HCP) (Sotiropoulos et al., 2013) and UK Biobank (Alfaro-Almagro et al., 2018) employ multi-shell dMRI as an essential imaging modality to depict the wiring of the human brain. Different from T1-weighted (T1w) and T2- weighted (T2w) MRI, which mostly capture anatomical details in the cerebral cortex and subcortical nucleus, dMRI provides rich information about the organization (Zeng et al., 2022) in white matter. The dMRI has provided many effective metrics to study whiter matter; for example, tensor-based scalar metrics, such as fractional anisotropy (FA) and mean diffusivity (MD)(Alexander et al., 2007), have been widely used for white matter characterization. More advanced modelling, such as the fibre orientation distribution (FOD) can be estimated with the sophisticated constraint spherical deconvolution (CSD) (Tournier et al., 2007)technique. Particularly, using multi-shell and high-angular resolution imaging protocols, CSD can better resolve crossing-fibre configurations in white matter, as well as observe more details of the neurite distribution in the cortical grey matter(Calamante et al., 2018). With the information from FODs, axonal streamlines with high angular resolution can be tracked with various algorithms(Calamante, 2019; Smith et al., 2020, 2012) and structural connectome(Yeh et al., 2019) can be constructed consequently. Recently, the “Fixel” (representing fibre population within a voxel) based analysis(Dhollander et al., 2021; Raffelt et al., 2017a) has been developed to quantitatively compare FODs in each voxel, and their derived features, such as apparent fibre density(Raffelt et al., 2017a) and fibre complexity(Dhollander et al., 2021; Raffelt et al., 2017a; Riffert et al., 2014), provide new means to quantitatively characterize the white matter.

In neuroimaging studies, a template space is essential for group-wise statistics and inference. A brain template is usually generated by averaging the spatially normalized brains from a representative population. Using it as a standard space, individual brains can be registered to the template for inter-subject comparison and group-wise statistical inference. For example, the Montreal Neurological Institute (MNI) T1w and T2w templates have served as the reference space for countless quantitative grey matter analyses (Ashburner and Friston, 2000) and functional analyses(Smith et al., 2004). A number of brain atlases are also defined in MNI space. Derived from diffusion tensor imaging, the MNI FA template has been used to guide white matter analysis(Evans et al., 2012), given its details in white matter. As the white matter has attracted increasing attention in neuroscience (Bethlehem et al., 2022; Zhu et al., 2013) and disease research, such as multiple sclerosis(Rovaris et al., 2005), Schizophrenia(Klauser et al., 2017; Lv et al., 2020) and Alzheimer’s Disease(Zhang et al., 2009), more quantitative and statistical analysis is desired on the complex FOD in white matter. A FOD template provides a reference space for such analysis.

The first FOD template was generated with the symmetric diffeomorphic registration by Rafelt et al (Raffelt et al., 2011). The template was generated solely with dMRI to minimize the inter-individual mean squared difference of FODs. With this FOD template as a reference, the Fixel Based Analysis (FBA) framework (Raffelt et al., 2017a, 2017b) was developed for group-wise inference. However, one limitation of this template is that the specificity of FODs at the vicinity of the grey-white matter boundary (peripheral white matter) is relatively low compared with that in the core white matter (Fig.2d). In these peripheral regions, however, the traditional registration approach based on T1w and T2w contrast provides better details for ensuring spatial specificity. A multi-modal approach for template construction therefore can offer substantial benefits(Greene et al., 2018; Uus et al., 2021).

In this paper, we propose to leverage the complementary information captured by T1w, T2w, and dMRI, exploring the classical tensor-based(Alexander et al., 2007) and the Fixel-based metrics(Dhollander et al., 2021), and use multimodal image registration(Avants et al., 2007) to generate a young adult brain template space, where the individual FODs are transformed into and averaged to be the FOD template. The HCP (Glasser et al., 2013) dataset is employed to guarantee the high quality of the template. As this template combines multimodal information, it contains good details for ensuring proper correspondence of both major white matter fibre bundles and grey matter regions, thus providing a tissue-unbiased FOD template. Anatomical constrained probabilistic fibre tracking(Smith et al., 2012) can be performed on this template to compute a tractogram template, and Tract-Seg (Wasserthal et al., 2018) can be performed to automatically segment the template into white matter bundles. As an illustration of the application of the new templates, the tractogram template is used to perform fibre-centred functional connectivity analysis(Lv et al., 2010), and group-wise functional connectivity is shown to be reproducible and stable.

## 2. Materials and Methods

### 2.1 Dataset

The HCP dataset(Glasser et al., 2013) was employed in this study, as it is among the best publicly available dMRI datasets currently. The acquisition parameters of the data are detailed below Please refer to (Glasser et al., 2013) for further details of the HCP acquisition protocol. dMRI was acquired using a spin-echo diffusion-weighted EPI sequence, with TR=5520ms, TE=89.5ms. FOV= 210×180mm^2^, matrix= 168×144, 111 slices, 1.25 mm isotropic resolution. Diffusion weighting consisted of 3 shells of b=1000, 2000, and 3000 s/mm^2^ and 90 directions per shell. Multiple b0 scans were collected throughout. A multiband factor of 3 was used to accelerate the acquisition(Glasser et al., 2013). Images were acquired using both phase-encoding polarities, for robust correction of susceptibility distortions.

HCP data also included high-quality T1w and T2w anatomical scans. The T1w data was collected using a 3D MPRAGE sequence, with TR=2.4s, TE=2.14ms, inversion time = 1s, flip angle of 8 degrees, FOV=224×224 mm^2^, 0.7 mm isotropic resolution. T2w data were collected using a 3D T2 SPACE sequence, with TR=3.3s TE=565ms, FOV=224×224 mm^2^, 0.7 mm isotropic resolution.

Resting-state fMRI (rfMRI) is another key imaging modality included in HCP for functional connectivity analysis. The rfMRI was collected using Gradient-echo EPI, with TR=0.72s, TE=33.1ms, flip angle of 52 degrees, FOV= 208×208 mm^2^, 2 mm isotropic resolution. A multiband factor of 8 was used to accelerate the scan. During the 14.5 minutes scan, 1200 volumes were collected.

Around 1200 young adults were recruited for the HCP project. In this study, we randomly selected 50 subjects (aged between 22 and 35, gender-balanced) for building the template. And we further randomly selected 300 participants for the fibre-centred functional connectivity analysis (see details in section 2.3).

### 2.2 Data processing

The minimally pre-processed data were downloaded from the HCP database, and we have performed additional processing steps of the dMRI data with the MRtrix3(Tournier et al., 2019) software. After performing the bias field correction(Tournier et al., 2019), the dMRI data were resampled to 1 mm to increase the quality of FOD. Two different dMRI models were considered for the template building. First, we calculated the tensor model (Alexander et al., 2007) and subsequently computed the FA and MD maps. Second, we used CSD (Tournier et al., 2007) to estimate the FODs. The tissue response function was estimated for each individual using the ‘Dhollander’ algorithm(Tournier et al., 2019) and then averaged over the 50 subjects to compute a group average response function. Multi-shell multi-tissue CSD (Jeurissen et al., 2014)was used to compute the FOD, accounting for partial volume effects. The intensity of white mater FOD was then normalized using the method in(Tournier et al., 2019). We used the first volume of the FOD (i.e., the l=0 term of the spherical harmonic expansion) to represent the total apparent fibre density (AFD_total_) (Raffelt et al., 2012b). And the FOD is further converted into Fixels, where the Fixel-based (Dhollander et al., 2021)feature ‘fibre complexity’ (Riffert et al., 2014) (CX) was estimated, which reflect the heterogeneity of fibre bundles. In summary, two parameters were computed from the tensor model (FA and MD), and two from the CSD model (CX and AFD_total_), which were used for the construction of the template.

The minimally pre-processed T1w and T2w images were downsampled to 1 mm isotropic in order to match the resolution of the dMRI. Additional pre-processing steps include spatial smoothing with FWHM=6mm and 0.01∼0.1HZ temporal bandpass filtering were applied to the minimally preprocessed resting-state fMRI data of HCP.

### 2.3 Template Generation Method

We employed the multivariate symmetric group-wise normalization (SyGN) (Avants et al., 2010, 2007) method to generate multimodal templates from the 50 HCP subjects. The SyGN robustly searches for a common space where each subject is transformed, and a template is generated by minimizing its difference with each subject in the group(Avants et al., 2010). The template generation can be summarized in the following steps(Avants et al., 2010):

1. Linearly align all the subjects and average them to initialize the template.
2. Pairwise normalisation between each individual and template with the symmetric normalization method(SyN)(Avants and Gee, 2004).
3. Average the registered individuals to update the template.
4. Go to 2) and repeat until the distance between the individuals and the template is minimized.

In our approach, we have used cross-correlation to define the energy function(Avants et al., 2008) for the optimization and the method converged after four iterations. The multivariate strategy (Avants et al., 2007)was used to combine the complementary information from the multiple imaging modalities and optimize the spatial normalization(Avants et al., 2007) and template optimization. In this paper, we have applied equal weights among different modalities. The following three approaches have been designed to generate the multimodal brain templates as shown in Fig.1a-c. Specifically, we used T1w, T2w images and feature images derived from dMRI as multi-channels for registration. The optimized transformation is used to warp the FOD into template space. Following reorientation(Raffelt et al., 2012a), the FOD images are averaged across subjects to generate the FOD template. Three different multimodal methods were designed in this paper. As illustrated in Fig.1, in *Method (a)*, we used only T1 and T2 images for template generation. This method is designed to test whether the registration based on the structural image alone can provide good FOD alignment. While in *Method (b)* we added tensor-based dMRI features of MD and FA. These well-recognized tensor metrics provide a better characterization of the white matter in healthy brains and diseased brains. In *Method (c)*, the FOD-based features of AFD_total_ and CX were combined with T1w and T2w to improve the template. AFD_total_ quantitatively measures the total neurite density within a voxel, while CX measures the heterogeneity of the fibres; they, therefore, also provide complementary information. These three templates are compared with the single-modal FOD-based Symmetric Diffeomorphic registration method(Raffelt et al., 2011) (Fig.1d), which is used as a baseline method given its widespread use for FBA and other dMRI analyses(Raffelt et al., 2011).

**Fig.1.**
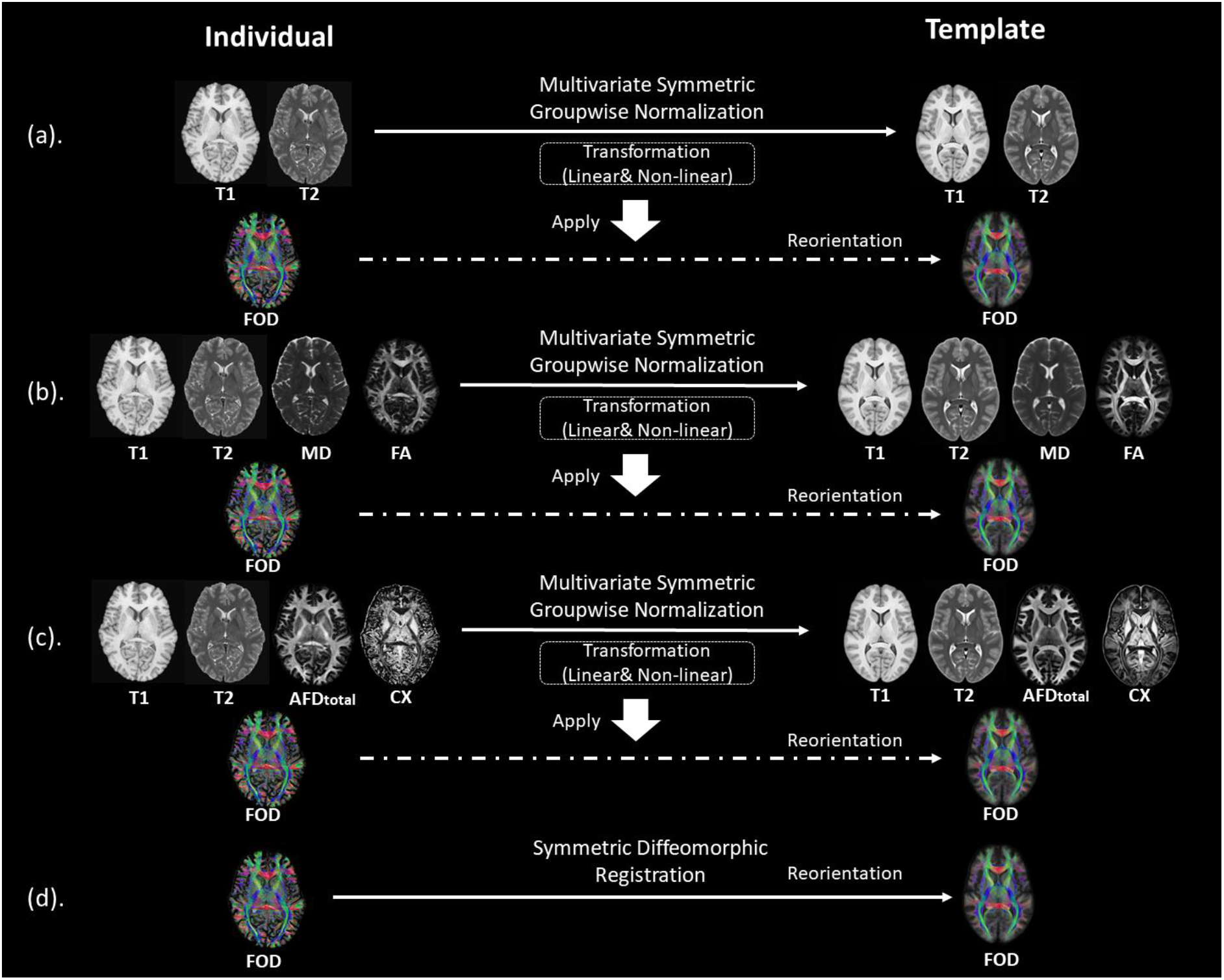
Method and Experiment design. (a-c) illustrates the multimodal registration and template generation. The transformation generated with multimodal images is applied to the FOD data, which are then averaged to generate each FOD template. (d) illustrates the population FOD template generation based on single modality FOD-based registration.

### 2.4 Evaluation

To evaluate the FOD templates, we calculated the average individual-template difference with two metrics: 1) root-mean-square error (RMSE) of spherical harmonics(Raffelt et al., 2011); and 2) Angular correlation coefficient (ACC) (Anderson, 2005). The two metrics reflect the intensity and angular alignment between individuals and the template, respectively.

We define *u*_*lm*_ and *v*_*lm*_ as the coefficients of the spherical harmonic expansion of the individual and the template FOD, respectively, at the corresponding voxel. land mrepresent the degree and order of a spherical harmonic function. The RMSE of the FOD is defined as:

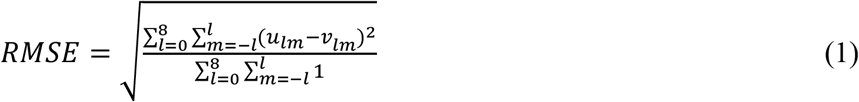

And ACC is defined as:

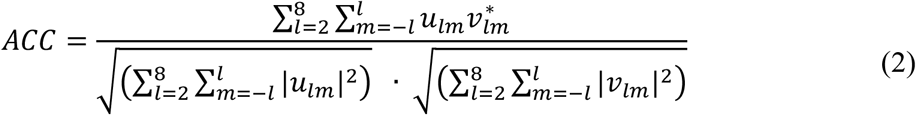

The evaluation was performed at two different scales, i.e., at the overall whole-brain white matter level and at the tract segmentation level. At the whole-brain white matter level, we averaged the individual-template difference of FOD of all the voxels within three brain masks defined based on various levels of AFD_total_. Specifically, the whole brain white matter mask is generated by thresholding with AFD_total_>0.1, the core white matter mask is defined with high total AFD (AFD_total_>0.2), and the peripheral white matter mask was defined with low total AFD (0.2>AFD_total_ >0.1) – see Fig.5. Within the three masks, we calculated the RMSE and ACC between the individual and template.

At the tract segmentation level, we first automatically segmented the whole brain into 72 tract bundles using the TractSeg toolbox(Wasserthal et al., 2018). The bundle masks and the bundle ending masks (Wasserthal et al., 2018) were compared between individuals and the template, and their agreement is measured by the Intersection over the Union (IoU) metric.

### 2.4 Generating the tractogram template with Anatomical Constrained Tractography

The multimodal template pipeline generates a set of templates in the input modality. As the T1w template generated with our method has the essential detail and good contrast between tissue types (see Fig.7a), we used Freesurfer(Fischl, 2012) to generate the tissue interfaces. In addition, the template tissue segments generated with the 5-tissue segmentation method (Smith et al., 2012; Tournier et al., 2019) using the T1w template can be used to guide the fibre tracking in the template, in a similar approach as it is used at the individual subject level fibre-tracking. To this end, we used the Anatomically Constrained Tractography (ACT) (Smith et al., 2012) framework using iFOD2 probabilistic fibre-tracking (Smith et al., 2012; Tournier et al., 2019)to generate the tractography template. The fibre tracking parameters were as follows: track number= 90,000, minimal length of 5mm and maximal length of 300mm. The other parameters are set with the default values provided by MRtrix3.

### 2.5 Calculating the fibre-centred functional connectivity

The fMRI data of HCP is warped to our template space to align with our tractography template. The image coordinates of the two endpoints were calculated (Lv et al., 2011, 2010)for each streamline of the tractogram template, and we then extracted the fMRI time series from those locations for each individual. The Pearson’s correlation of the fMRI signals at the two ends was used to represent the functional connectivity associated with the streamline(Lv et al., 2011, 2010; Zhu et al., 2014).

## 3. Results

We present below an evaluation of the proposed FOD templates in comparison with the existing baseline method, as well as demonstrate a possible application with the unbiased template, namely fibre-centred functional connectivity using the tractography template.

### 3.1 Whole-brain **FOD Templates**

With the four experimental settings introduced in Section 2.3, we have generated four sets of FOD templates. We qualitatively show a cross-section of the four FOD templates in Fig.2, overlayed on the AFD_total_ (i.e. *l*_0_ term of the FOD). FOD templates generated with the multimodal methods (Fig.2a-c) clearly preserve the gyrus and sulcus organisation and provide much higher spatial specificity near the white-grey-matter boundaries than the FOD-based template (Fig.2d). As can be observed, the multimodal methods also perform similarly well at crossing fibre zones. We further visualized the AFD_total_ from multiple perspectives in Fig.3. It is evident that the multimodal methods deliver sharp contrast (see the first 3 rows in Fig.3) at the white-grey-matter boundaries, in the cerebellum, in subcortical structures (such as thalamus and hippocampus), and preserve the cortical folding patterns in detail (see the volume rendering at the fourth row of Fig.3).

**Fig.2.**
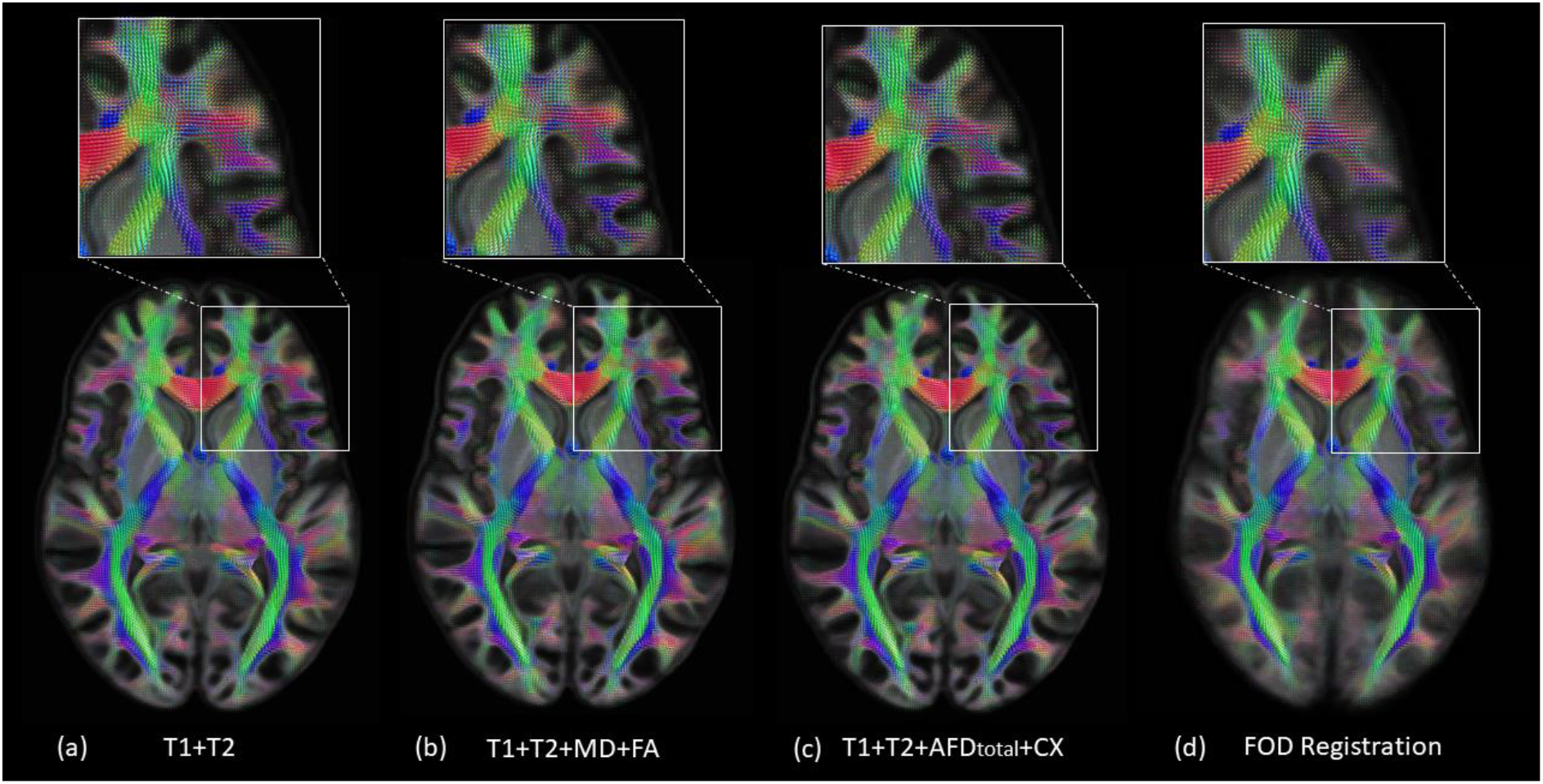
FOD templates generated from 50 HCP subjects with the 4 strategies described in Fig. 1. The FOD butterfly patterns are overlaid on the AFD_total_ map, with the colours indicating the local fibre orientation (red: left-right; green: anterior-posterior; blue: inferior-superior). The white boxes highlight zoomed areas of the FODs with crossing-fibres and near-cortex white matter.

**Fig.3.**
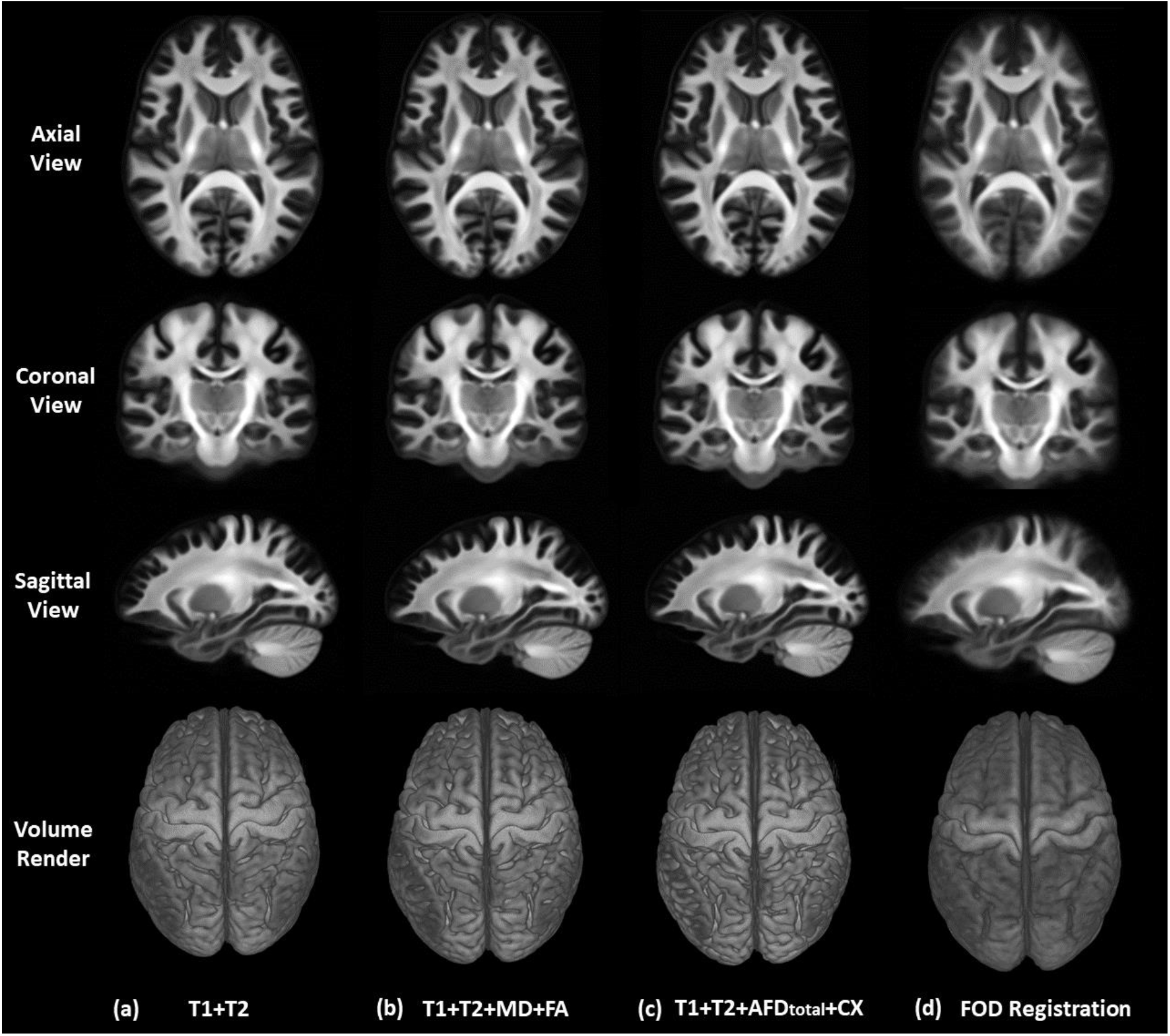
Visualization of AFD_total_ map from each of the four FOD templates. The rows represent three viewing perspectives and the volume rendering. The result of the traditional FOD-based registration, which is shown in (d), has blurry white-grey-matter boundaries, and ambiguous folding patterns. In contrast, the results of our multimodal methods in (a-c) are much sharper.

The two individual-template difference metrics (RMSE and ACC) were averaged across 50 subjects and visualized in Fig.4. Qualitatively, we can see that our method shows lower RMSE and higher ACC near the grey matter regions compared with the FOD-based registration method. For each subject, we further calculated the overall individual-template difference by averaging each metric across voxels within the brain masks defined in Section 2.3. The averaged differences and the standard deviation based on the statistics of 50 subjects are plotted in Fig.5 for the whole brain white matter, the core white matter (with high AFD_total_), and the peripheral white matter (with low AFD_total_). As shown in Fig.5b, within the whole brain white matter mask, both the T1+T2+MD+FA and the T1+T2+AFD_total_+CX methods show the lowest RMSE of FOD and highest ACC (p<0.05, FDR), while the T1+T2+MD+FA method performs slightly better than T1+T2 + AFD_total_+CX in terms of ACC. At the core white matter regions (Fig.5c), both the T1+T2+MD+FA and the T1+T2+AFD_total_+CX methods show comparable RMSE with FOD registration and the T1+T2+MD+FA show slightly better (p<0.05, FDR) ACC than the FOD-based registration. The largest difference takes place at the peripheral white matter(Fig.5d), where both the T1+T2+MD+FA and the T1+T2+AFD_total_+CX methods show much lower RMSE (p<0.05, FDR) and much higher ACC (p<0.05, FDR) compared with FOD-based registration. This indicates that multimodal registration is capable of generating a tissue unbiased FOD template. As a reference, the T1+T2 method showed relatively good performance at the peripheral white matter but poor performance at the core white matter regions, which is reasonable because both T1w and T2w images provide limited detail in the white matter.

**Fig.4.**
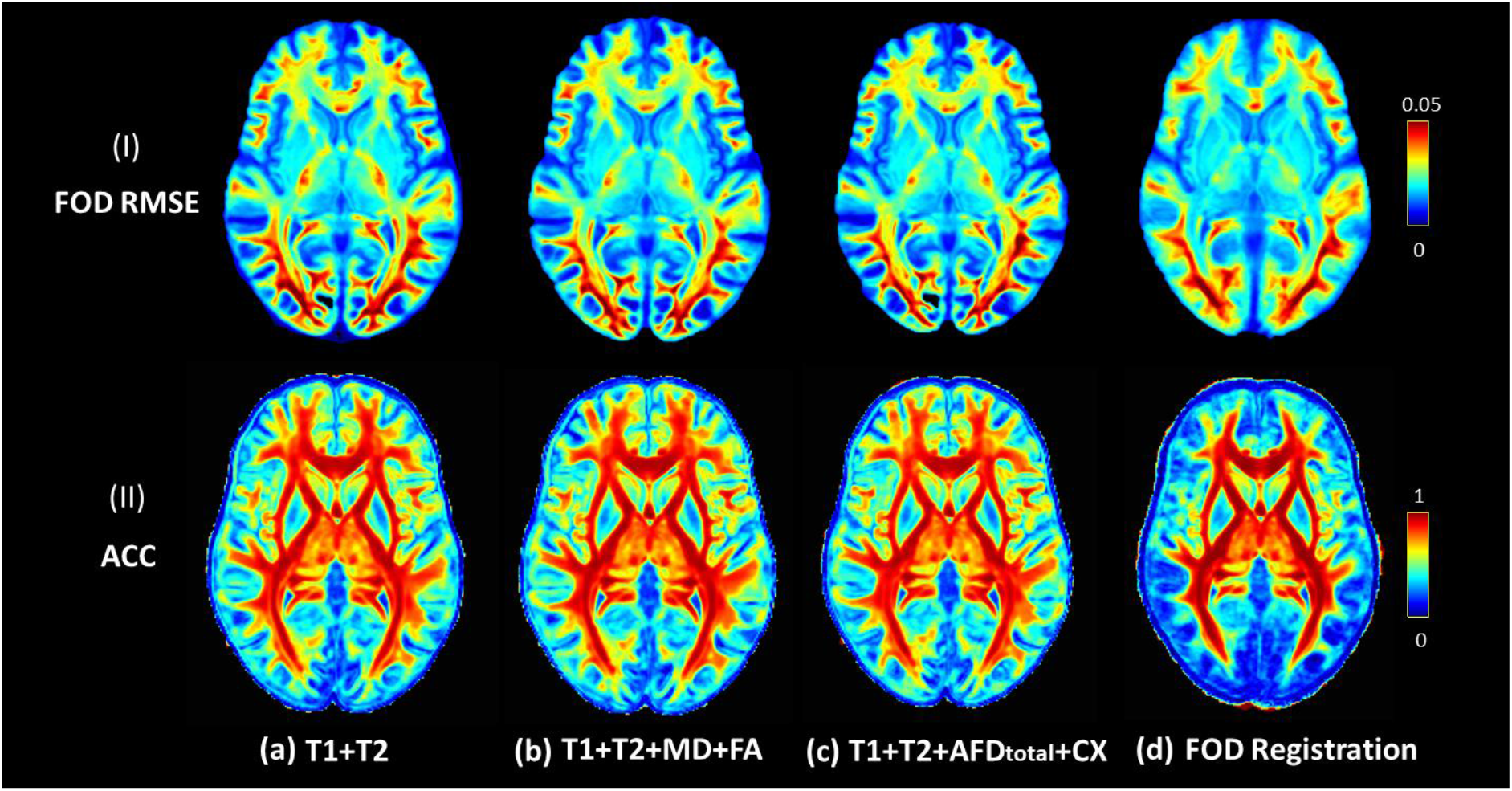
The average individual-template difference calculated by averaging the metrics across subjects. Each row shows the averaged voxel-wise measurement across 50 subjects. The lower RMSE of FOD indicates better amplitude agreement, while the higher angular correlation indicates better angular alignment.

**Fig.5.**
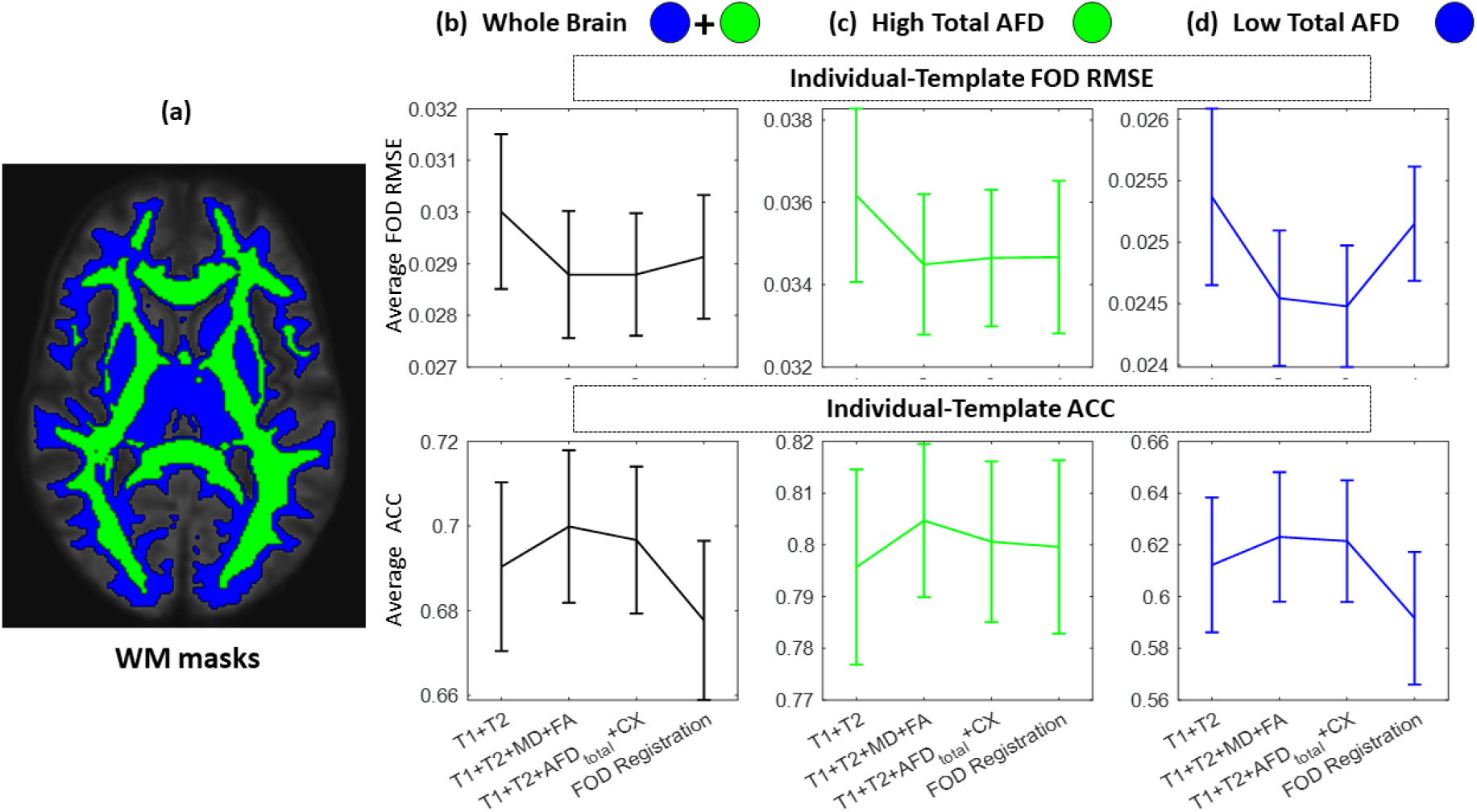
The individual-template difference calculated by averaging each metric across all the voxels within different white matter(WM) masks (a). The error bar plots show the mean and standard deviation of the 50 subjects. The individual template-difference are compared between the four methods within the whole brain WM mask (b), core WM mask(c), i.e. with high total AFD (AFD_total_>0.2), and peripheral WM mask(d) with low total AFD (0.2>AFD_total_ >0.1). The first row shows the difference in terms of the average RMSE of FOD, while the second row shows the difference in terms of average ACC. Similar to Fig.4, The lower overall RMSE of FOD indicates better amplitude agreement, and the higher ACC indicates better angular alignment.

### 3.1 Comparison of the Tract Segmentation

The Tractseg method automatically produces 72 white matter tract segmentation, which has been widely used to evaluate the major fibre bundles in the brain. Fig.6a shows 2 example fibre bundles: the right Corticospinal tract (CST) and right Inferior occipitofrontal fascicle (IFO). Each bundles’ spatial geometry can be characterized by two segmentation masks, the bundle segmentation and the ending segmentation. We have applied Tract-Seg separately on the 4 FOD templates, as well as on the corresponding individual FODs registered in each template space. The IoU index (for which higher values indicate better performance) was used to measure the spatial individual-template agreement of the 72 bundle and 144 ending segmentations. The histogram of the bundle(Wasserthal et al., 2018) IoU and the ending segmentation(Wasserthal et al., 2018) IoU are shown in Fig. 6b and 6c, respectively. And the outline of each histogram (with colour correspondence) shows the fitting of the distribution. Overall, both the T1+T2+MD+FA and the T1+T2+AFD_total_+CX methods show the best agreement (high IoU) between individual and template on the 72 fibre bundles.

**Fig.6.**
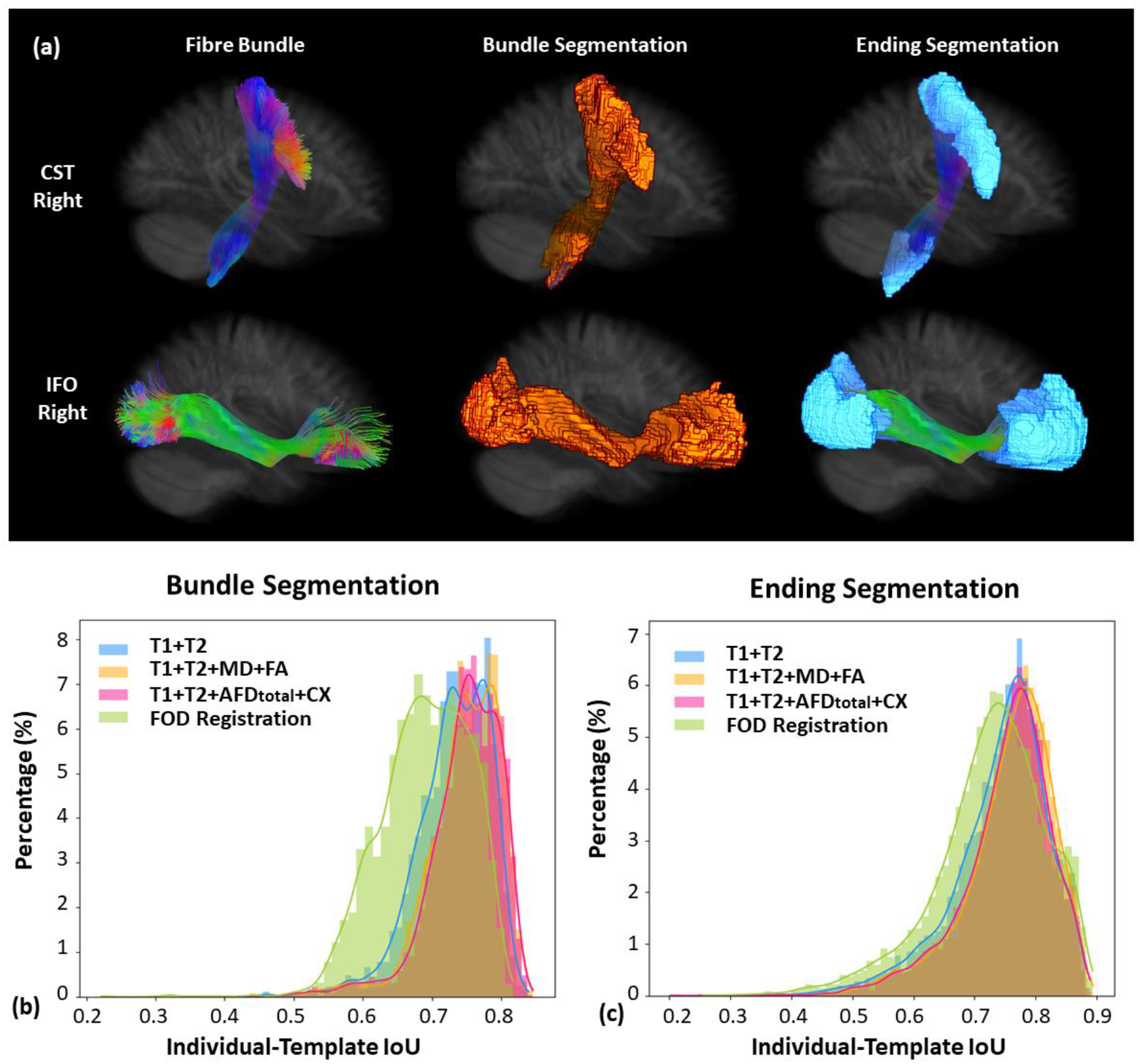
The histogram of IoU index of the tract segmentations between individual and templates. (a) Two examples of the 72 fibre bundles (right Corticospinal tract (CST) and right Inferior occipitofrontal fascicle (IFO)) generated by Tract-Seg. (b) The percentage histogram of IoU of bundle segmentation. (c) The percentage histogram of the IoU of the ending segmentations.

### 3.2 Anatomical Constrained Tractography Template

Compared with the traditional FOD-based registration method, the multimodal methods in this paper also produce templates of each of the input modalities, including structural T1w and T2w templates. As shown in Fig.7a, we have visualized the T1w template generated by the T1+T2+MD+FA method. This template provides fine details in the cortical regions, subcortical regions and the white-gray-matter boundaries. Qualitatively, this template looks much finer than the MNI T1w template. As a result, we have successfully reconstructed the cortex surfaces with fine anatomical detail, as shown in Fig.7g. Similarly, we can also generate the five tissue probability maps with the 5-tissue segmentation method(Tournier et al., 2019), corresponding to cortical gray matter, subcortical gray matter, white matter, cerebrospinal fluid, and abnormal tissue (which is not present in our dataset of healthy subjects). The results can be seen in Fig.7b- e.

**Fig.7.**
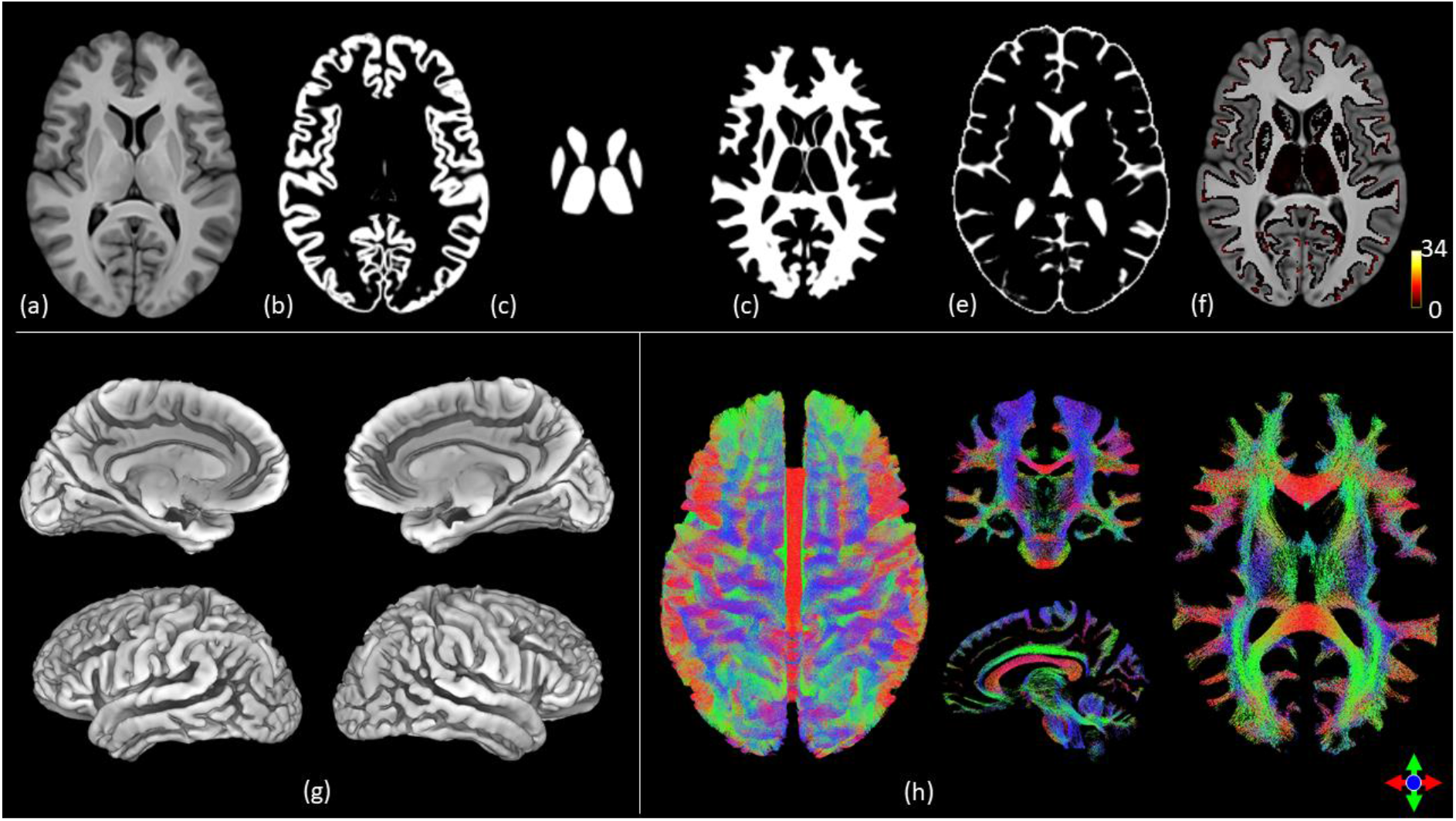
(a) The T1 template generated from the T1+T2+MD+FA method. (b-e). The cortex, subcortex, white matter and CSF regions generated by the 5ttgen method within MRtrix based on the T1 template in (a). (f) The fibre ends of the template tractogram (h) overlayed onto the T1 template. (g) The white-gray matter interface constructed by Freesurfer using the T1 template in (a). (h) The fibre tracks of the template tractogram generated by ACT method using the FOD template generated by the T1+T2+MD+FA method, displayed as a view from above, or as slices in three orientations (coronal, sagittal and axial). The colours in (h) indicate the local fibre orientation (red: left-right; green: anterior-posterior; blue: inferior-superior).

With the five tissue maps generated, we also successfully performed fibre tracking using the Anatomically Constrained Tractography framework(Smith et al., 2012). The generated tractogram template is visualized in Fig.7h. Qualitatively, the streamlines reflect the known white matter anatomy. Particularly, the template nicely characterises areas of crossing fibres. We have further mapped the fibre ending density and overlaid it on the T1w template. As shown in Fig.7f, the fibre ends densely distributed throughout the gray-white-matter boundaries.

### 3.3 Fiber-Centred Functional Connectivity

The tractogram template generated in section 3.2 has high specificity at the white-grey-matter boundaries and therefore provides a good substrate to calculate the fibre-centred functional connectivity(Lv et al., 2011, 2010) of each individual with their rsfMRI data warped to the template space. The derived functional connectivity reflects the strength of functional synchrony between the two fibre endpoints. We have colour coded the functional connectivity onto the streamlines for visualization. In these images, reddish (blueish) pathways indicate fibre bundles that connect gray matter regions with high (low) functional synchrony. To evaluate the reliability of the fibre-centred functional connectivity, we have randomly divided the 300 HCP subjects into 3 group, 100 subjects in each group without overlap. We have averaged fibre-centred functional connectivity within each group, and these results are visualized in Fig.8. High consistency of functional connectivity (with spatial correlation of 0.9939±0,0002, p<10^−10^) is seen across the average of three sets of 100 subjects(Fig.8).

**Fig.8.**
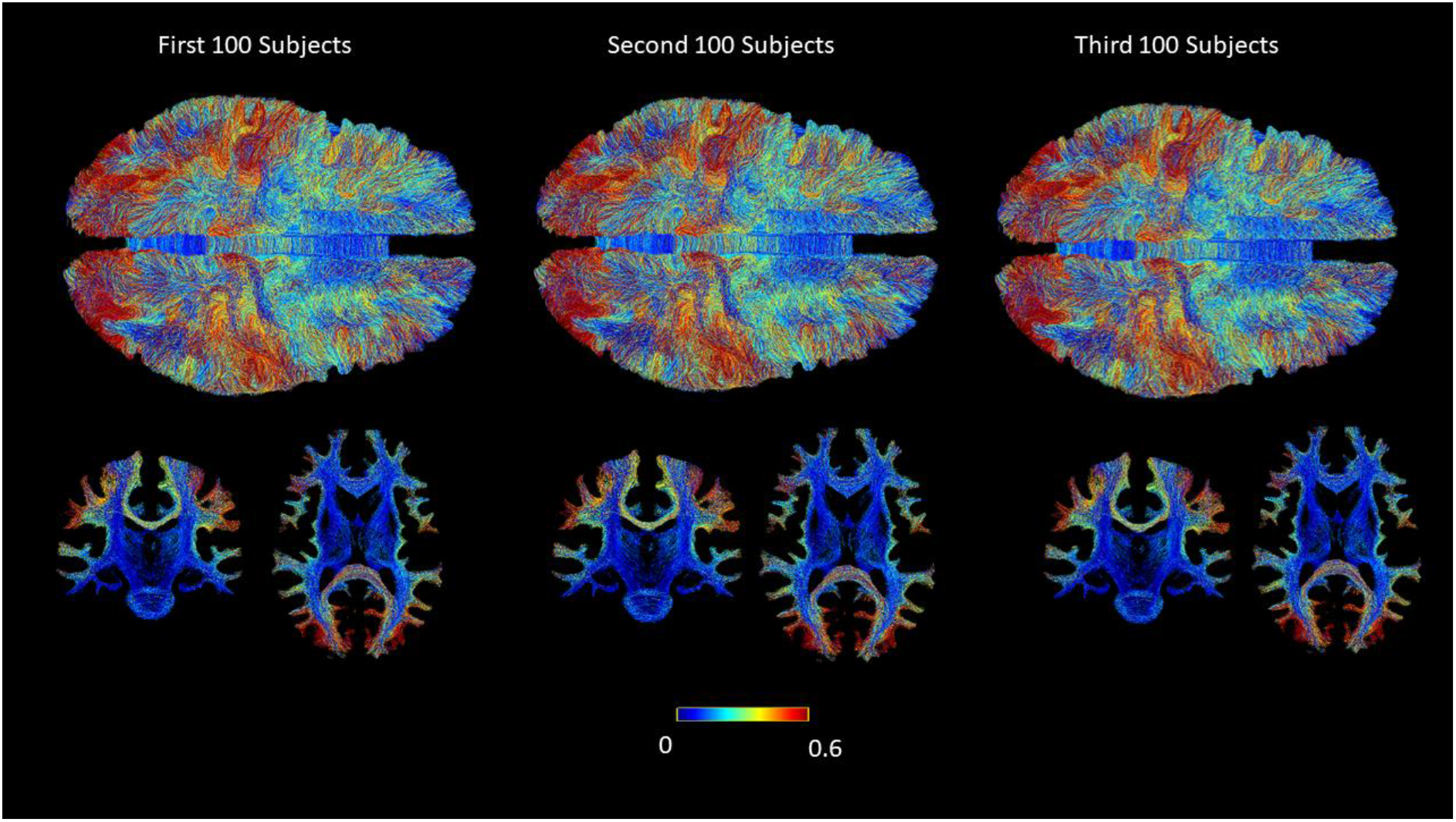
Fiber-centred functional connectivity results. The functional connectivity measured by the Pearson’s correlation of two endpoints of the streamline mapped (as colour) onto the streamline. The three columns are the average results from three sets of subjects (100 subjects each). The top row shows the results view from above; the bottom row shows coronal and axial sections.

## 4. Discussion

We have provided a new framework to build a white matter FOD template using multimodal MRI data. Multimodal registration allows combining the complementary information from the T1w and T2w (with high cortical and subcortical gray matter detail) and Diffusion MRI (with high white matter detail), and thus generate a tissue-unbiased template space. The FOD templates generated by combining T1w, T2w and Diffusion MRI features have shown high detail in both core white matter regions and peripheral white matter regions, even for subcortical white matter where the traditional FOD-based template lacks specificity. The associated T1w and T2w templates, which are determined during the process of multimodal template construction, also provide very fine detail for the grey matter and grey-white-matter boundaries. We thus conclude that the approach proposed generates a robust tissue-unbiased FOD template for neuroscience applications.

The conventional FOD template generated with the FOD-based registration(Raffelt et al., 2011) has shown very good specificity on core white matter regions but compromised specificity at the peripheral white matter and grey-white-matter boundaries. This has caused difficulties for joint grey and white matter analysis or joint structural and functional analysis, as traditional templates tend to favour one tissue type at the expense of the other. Therefore, the tissue-unbiased FOD templates and the associated tractogram template generated in this paper has filled an important gap in the field. For example, with our tractogram template, fibre activation detection method proposed in (Lv et al., 2011) can be now evaluated at the group level.

As shown in Fig.7, the multimodal T1w(T2w) template presents abundant details for tissue segmentation and surface reconstruction. In contrast, when we run the Freesurfer and 5-tissue segmentation on the widely used MNI T1w template, both methods fail to generate a result; this is likely because of the poor grey matter and white matter contrast and ambiguous white-grey-matter boundaries in the MNI template. On the other end, the FOD template generated by solely FOD registration cannot provide a brain structure template for tissue segmentation. This stresses an advantage of the proposed multimodal template approach. Only with this advantage, the ACT method can be used to generate the tractogram template with nice alignment with the grey-white matter boundary. Consequently, our method provides new potential for joint grey-white matter analysis and joint structure-function analysis.

One well-recognized challenge for neuroimage analysis is the lack of ground truth. This also applies to the building of brain templates. To evaluate the quality of the FOD template, we have proposed to use the strategy of individual-template difference. We believe it is a fair comparison so far in the field. For the fibre-centred functional connectivity, we have also cross-validated the results across separated groups. The consistency across the groups has shown the strong reliability of the analysis.

In this paper, we have explored only a limited number of combinations of diffusion MRI metrics to combine with T1w and T2w structural images for template generation. The results have been very promising. More importantly, we have provided a new framework for building tissue-unbiased FOD templates from multimodal MRI, which could be easily extended to incorporate alternative diffusion MRI metrics.

The FOD template is generated by averaging the registered individual FODs. The fibre tracking algorithm can track the common tracks with the FOD template, but inevitably it may generate spurious fibres. We did not cover the validation and refinement of the tractography template. In our future work, we will explore the benefits of fibre filtering methods, such as spherical-deconvolution informed filtering (SIFT)(Smith et al., 2013) and SIFT2(Smith et al., 2015) methods, for refinement of the tractogram template.

The axonal fibres are the substrate of regional functional connectivity. More and more interest has been shown to the axonal connection informed the functional connectivity (Calamante, 2017; Calamante et al., 2017, 2013; Lv et al., 2011, 2010). In this paper, we have provided a new method to define fibre-centred functional connectivity, which reflects structural-informed functional synchrony(Calamante et al., 2013; Lv et al., 2010). With the tractogram template as a reference space, group-wise and cross-group analysis can be performed in the future, providing a new framework for investigating structural-functional properties in neuroscience and clinical applications. Particularly, our template can be used to enhance the fixel-based analysis(Dhollander et al., 2021) at the gray-white matter boundaries and the peripheral white matter (close to gray matter) to detect anomaly. The tractogram template itself is a high-resolution connectome(Mansour L et al., 2021), which reflects the commonality of the human brain. This is different from the traditional region-region connectome, but a fibre-centred connectome. As a reference system, the fractogram template provides a new mean to aggregate features from multiple subjects and datasets for machine learning approaches(Ganesan et al., 2021; Lv et al., 2017, 2015).

## 5. Conclusion

We have proposed a new methodology framework for building a tissue-unbiased FOD template based on multimodal registration. Based on the resulting FOD template, a tractography template using anatomical constrain was demonstrated. It has provided a new potential for joint grey matter and white matter analysis, and joint structure and function analysis.

## Supporting information

The 3D visulization of the tractogram template.

## 6. Acknowledgement

We acknowledge the data made available through the Human Connectome Project, WU-Minn Consortium (Principal Investigators: David Van Essen and Kamil Ugurbil; 1U54MH091657) funded by the 16 NIH Institutes and Centers that support the NIH Blueprint for Neuroscience Research; and by the McDonnell Center for Systems Neuroscience at Washington University. We acknowledge the support by the MASSIVE HPC facility (www.massive.org.au). FC is supported by the NHMRC (#APP1117724) and ARC (#DP170101815).

## Credit author statement

JL and FC conceptualized the study.

JL, RZ, MH and AD processed and analysed the imaging data.

JL and FC wrote the initial draft.

All co-authors edited the final manuscript.

Declarations of conflict of interest: None.

